# Unraveling the diversity of protein-carbohydrate interfaces: insights from a multi-scale study

**DOI:** 10.1101/2024.10.22.619630

**Authors:** Aria Gheeraert, Frédéric Guyon, Serge Pérez, Tatiana Galochkina

## Abstract

Protein-carbohydrate interactions play a crucial role in numerous fundamental biological processes. Thus, description and comparison of the carbohydrate binding site (CBS) architecture is of great importance for understanding of the underlying biological mechanisms. However, traditional approaches for carbohydrate-binding protein analysis and annotation rely primarily on the sequence-based methods applied to specific protein classes. The recently released DIONYSUS database aims to fill this gap by providing tools for CBS comparison at different levels: both in terms of protein properties and classification, as well as in terms of atomistic CBS organization. In the current study, we explore DIONYSUS content using a combination of the suggested approaches in order to evaluate the diversity of the currently resolved non-covalent protein-carbohydrate interfaces at different scales. Notably, our analysis reveals evolutionary convergence of CBS in proteins with distinct folds and coming from organisms across different kingdoms of life. Furthermore, we demonstrate that a CBS structure based approach has the potential to facilitate functional annotation for the proteins with missing information in the existing databases. In particular, it provides reliable information for numerous carbohydrate-binding proteins from rapidly evolving organisms, whose analysis is particularly challenging for classical sequence-based methods.

## INTRODUCTION

Carbohydrates are fundamental organic compounds in biological and biochemical pathways across diverse ecosystems. Interactions between proteins from various functional classes and carbohydrates are essential, as they maintain life, holistic tissues and homeostasis, thereby governing a wide range of biological processes from energy generation^1^ to cellular recognition^2–5^. Protein-carbohydrate interactions involve cell proliferation, differentiation, aggregation, signal transduction, inflammation, host-pathogen recognition^6–9^ and protein structure stabilization. As such, they have numerous uses in the design of pharmaceuticals^10,11^, including creating antibodies, vaccinations, and inhibitors. Understanding the underlying mechanisms at the molecular level requires a detailed description of protein-carbohydrate interface diversity at different scales, from the macroscopic protein properties to the structure and composition of binding sites.

Despite the advances in annotation and classification of carbohydrates and carbohydrate-binding proteins^12–25^, information on protein-carbohydrate interaction patterns remains sparse. First, the number of experimentally resolved protein-carbohydrate complexes is limited due to the inherent flexibility of these interfaces. Secondly, general databases such as Protein Data Bank^26^ (PDB) lack coherence in annotating carbohydrates due to their chemical diversity. Indeed, they can comprise one or more monosaccharide units, have linear or branching structure, and exist freely or bound to other molecules. Across this article, we will use the term “sugar” or “carbohydrate” to denote monosaccharides, oligosaccharides, polysaccharides, and glycoconjugates. Thirdly, while information on sequences and available structures of carbohydrate-binding proteins can be found in different specialized databases, those are maintained by different research groups, do not focus on the same type of annotations and do not contain information on the carbohydrate binding site properties. Carbohydrate-Active enZYme (CAZy) database^20,27^ classifies proteins with catalytic activity involved in different stages of glycan synthesis, modification, and degradation. These include the glycosyltransferases, the glycoside hydrolases, the carbohydrate esterases, and proteins involved in synthesizing and transporting precursor molecules for glycosylation reactions, including nucleotide-sugars. CAZy also contains annotation of the proteins binding carbohydrates without altering the ligand, such as carbohydrate-binding modules (CBMs). The reference source of information on lectins is UniLectin^13^, containing both annotation and classification of lectins, as well as their structures^12^ and tools for sequence-based discovery of new lectins^14^. 3D structures of protein complexes with glycosaminoglycans (GAG) for six most common mammalian GAGs are available in GAG-DB^28^. Finally, information on glycan-antibody interactions can be extracted from the SAbDab database of annotated antibody structures^19^. The general database of the resolved structures of protein-carbohydrate complexes, ProCarbDB^16^ was published in 2021 but is unfortunately no longer available. Finally, carbohydrate binding sites (CBS) are implicitly included to the protein-ligand interaction databases such as BioLiP^29^ or Binding-MOAD^30^. However, such databases often miss out some essential carbohydrate properties by mixing up covalent and non-covalent interactions.

In 2024 our group has for the first time retrieved and annotated all carbohydrate-containing entries available in the PDB and gathered all the relevant information available in the existing databases in the form of a free open-source platform DIONYSUS (https://www.dsimb.inserm.fr/DIONYSUS/)^31^. In the current study, we provide an in-depth exploration of DIONYSUS content with focus on the non-covalent protein-carbohydrate interactions. We analyze carbohydrate and protein diversity in terms of sequence, fold and function, as well as at the level of local geometrical resemblance among the interfaces formed by proteins and carbohydrates. Such multi-level comparison allowed us to address the following questions: what is the actual amount of different protein-carbohydrate interfaces in the PDB? What is the diversity of carbohydrate binding sites for similar proteins? How can we evaluate the redundancy of CBS data for different protein classes? Furthermore, our rigorous annotation discovered similarities between carbohydrate binding sites in seemingly evolutionarily unrelated folds.

Finally, the suggested method of CBS comparison coupled to the exhaustive nature of our database allowed us to provide high-confidence annotations for tens of lectins absent from UniLectin and several carbohydrate binding modules with no CAZy annotation, therefore completing traditional sequence-based predictions. Notably, our methodology efficiently discovers CBS in proteins coming from organisms prone to substantial sequence alterations, such as bacterial and viral lectins and binding modules from human gut microbiota.

## MATERIALS & METHODS

In the framework of our study we focus on four functional classes of carbohydrate binding proteins: lectins, carbohydrate active enzymes, carbohydrate binding modules and antibodies. These protein classes are the most abundantly present in complex carbohydrate ligands in the PDB. We discuss further extension of our methodology to other important functional classes of carbohydrate-binding proteins in Discussion.

All the datasets used in this study were extracted from the DIONYSUS database^31^, version of June 2024. Comparison of the protein-carbohydrate interface geometry was performed using a graph-based score implemented in DIONYSUS compare tool (www.dsimb.inserm.fr/DIONYSUS/compare.html) with source code available at: www.github.com/DSIMB/CompareCBS. Optimal alignment between two binding sites is calculated from a mapping between the atoms of the same type providing a minimal possible distortion of the interatomic distances^32–34^. The resulting score varies between 0 and 1, where 1 means a perfect match between binding site atoms with exactly the same internal distances and 0 implies that all internal distances between the same pairs of atoms are above a given precision threshold (1Å in our case).

Clusters of similar CBS identified in DIONYSUS^31^ were further explored regarding the available annotations in specialized databases such as CAZy^20,27^ and UniLectin^12^. Information on organisms and the kingdom of life was extracted from UniProt. Protein fold type was attributed according to Evolutionary Classification of PrOtein Domains (ECOD^35^). Sequence identity was evaluated using MMseqs2^36^. Protein structural similarity was computed using TM-score^37^.

## RESULTS

### The actual number of different protein-carbohydrate complexes in the PDB

#### Carbohydrate diversity of the PDB

According to DIONYSUS annotations, almost 13% of the ligands catalogued in the PDB possess a ring with sugar-like properties. Indeed, we retrieve >3k different chemical components containing carbohydrate residues (found in mono-, oligo- and poly-saccharides, and in glycoconjugates, Fig. S1-S2). For comparison, only 971 components are currently annotated as saccharides in PDB^38^, among which 833 are identified as non-obsolete and cyclic^21^, and 708 carbohydrate residues were reported in ProCarbDB^16^. Our exhaustive annotation of carbohydrates allows us to retrieve >47k PDB entries containing protein-carbohydrate complexes (Fig. 1a), which is three times more than reported in previous studies (13k in ProCarbDB^16^, and 14k in the recent study^21^).

**Figure 1.**
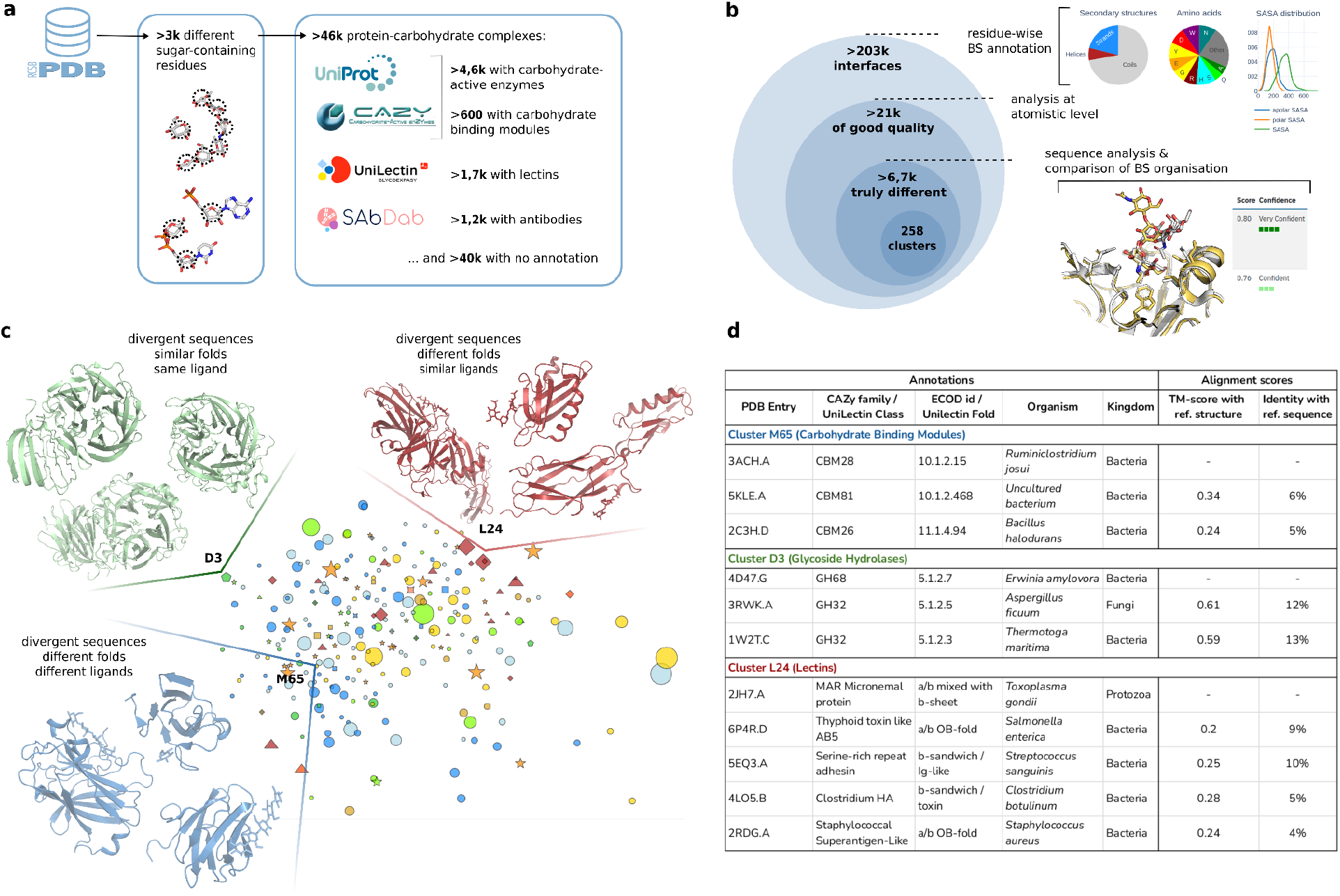
**a**. Data extraction and annotation according to different specialized databases. **b**. A number of non-covalent interfaces formed by carbohydrates according to different criteria. Detailed information is provided in the Supplementary Information section “Diversity of carbohydrate binding sites for different proteins” (Fig. S3 and Fig. S4). **c**. t-SNE projection of the clusters of similar carbohydrate binding patterns. Each cluster is represented by a SNFG symbol of the most frequent carbohydrate. **d**. Detailed information on the most divergent proteins found in clusters highlighted in panel c. Colors of the cluster names correspond to those from panel c. Structural diversity of proteins is evaluated using the Evolutionary Classification of PrOtein Domains (ECOD^35^).

#### Carbohydrate-binding protein diversity of the PDB

Despite the constantly developing specialized databases of protein-carbohydrate interactions, almost two-thirds of the extracted binding sites are currently not annotated in the considered databases (Fig. 1a). Among them, 28 different GAG-binding proteins are annotated in GAG-DB^28^. Among the remaining proteins, according to the analysis of the PDB keywords, about 63% of these binding sites correspond to enzyme CBS (PDB keywords: “hydrolase”, “transferase”, “oxidoreductase”, “lyase”, or “isomerase”) but not annotated as active site by UniProt. Over 12% of diverse CBS among “Others” contain lectin-associated keywords in their annotation, such as “Sugar binding protein”, “viral protein”, “metal binding protein”, “cell adhesion”, but the corresponding proteins are not referenced in UniLectin (Fig. S4-S5). As shown below in the “Filling missing annotations” section, most of the corresponding sites can be attributed to one of the known lectin families. Finally, proteins with completely different functions are also referenced in our database, which is the most common class corresponding to transport proteins (4% of unannotated sites). Immune system proteins, which are not referenced in SabDab (e.g., PDB ID 6IFJ), only represent 1% of the “Others” binding sites.

#### Carbohydrate binding site diversity of the PDB

Not all interfaces reported in DIONYSUS are of the same quality and biological relevance. Indeed, experimental difficulties in carbohydrate resolution lead to the presence of numerous structures with missing atoms or clashes. Moreover, carbohydrates are frequently used as crystallographic adjuvants and therefore, can form complexes, which are never observed in biological systems. Only 11% of 203k non-covalent interactions are resolved without artifacts and are referenced as biologically relevant according to BioLip^29,39^ and were further used to identify representative binding patterns (Fig. 1b). Nevertheless, in depth comparison of the selected 21k binding sites reveals that only 6,7k of them can be considered as truly different, while others are formed by the same protein and same carbohydrate at exactly the same regions on the protein surface. We have also identified several cases of identical proteins binding the same carbohydrate residue forming different patterns (Fig. S5).

Finally, the pairwise comparison of CBS geometry^31^ identified 258 clusters of most common patterns of protein-carbohydrate interactions: 79 clusters of CAZy active sites (68 **D**egradation, 3 **S**ynthesis, 7 **P**olysaccharide Lyase and 1 au**X**iliary activity), 84 clusters of lectin-carbohydrate interactions, 91 clusters of CBM binding sites and 4 clusters of antibody binding sites (Fig. 1c).

### Revealing carbohydrate binding site convergence in proteins from different families

Analysis of three-dimensional organization of carbohydrate binding sites allowed the detection of characteristic binding patterns, which can be found in proteins with extremely divergent sequences and/or structures. The most prominent examples of such convergence are shown in Fig. 1c.

#### Convergence of CAZy and CBM binding sites

Several clusters of glycosyl hydrolases gather proteins from various families responsible for the same function (Tab. S1). For instance, three glycoside hydrolases from cluster D3 (Fig. 1c, in green) share a very similar binding site responsible for fructan degradation, but belong to different CAZy families, are found in different kingdoms of life and share very low sequence identity (Fig. **1d**, in green).

Binding site convergence for proteins of different folds is even more for carbohydrate binding modules. For example, cluster M65 gathers bacterial CBMs attributed to different families (Fig. **1d**, in blue) but forming very similar interfaces with glucose moieties in both mono- or oligosaccharide forms. Moreover, in general, CBM binding sites form smaller and more diverse clusters than enzymatic ones (Tab. S3, Fig. S6-S10), which is explained by weaker evolutionary pressure as compared to the enzymatic active site regions. However, thorough investigation is hindered by lack of consistent annotations, a big part of which was finally done manually (see Tab. S2 for further details).

#### Sialic acid binding patterns in pathogens and animal lectins

Binding site conservation among different species and folds is also characteristic for lectins. We detect eleven clusters of lectin binding sites containing proteins from different Unilectin classes (Tab. S5). One of the most diverse of them is L24 (Fig. 1c, in red; Fig. 2a) gathering sialic acid binding sites found in pathogens from bacteria or protozoa. These binding sites are responsible for recognition of α2,3- or α2,6-sialylated ligands on cell surfaces during host-pathogen recognition. Our method reveals their almost identical organization on the protein surface (Fig. 2a, zoom) for proteins of extremely divergent sequences and folds (Fig. 1d, red). Indeed, in all CBS of cluster L24 sialic acid residue is oriented in the same way in respect to the β-sheets in CBS composition and the corresponding β-sheet region has the same composition for all the observed structures (Fig. 2b). Interestingly, sialic acid extends the hydrogen bond pattern of the parallel β-sheets using their amide and carboxylic acid moieties, mimicking another β-sheet (Fig. 2b, Fig. S11).

**Figure 2.**
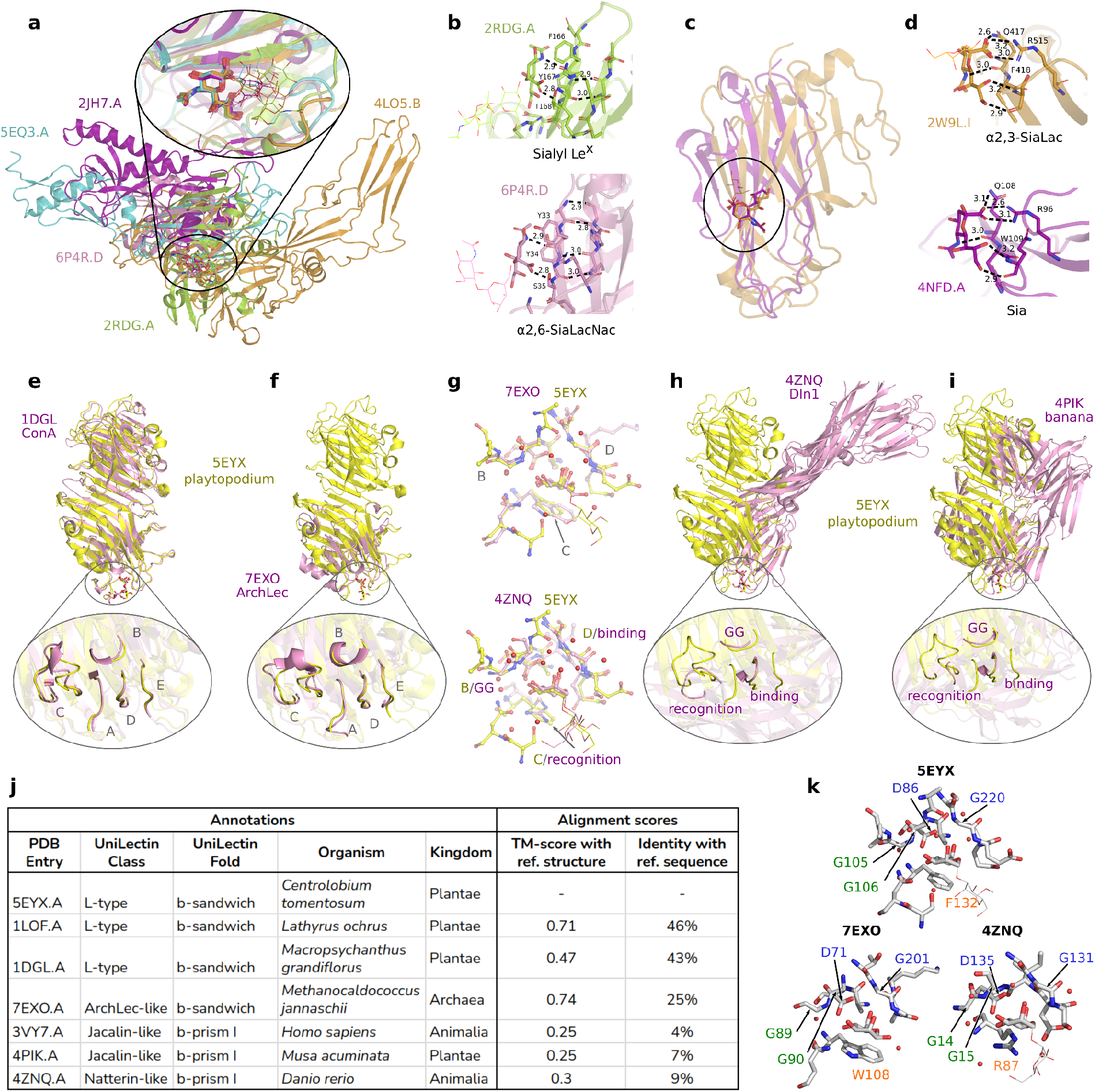
Different clusters of lectin binding sites. **(a-d)** Superposition of protein structures from clusters L24 (**a**) and L25 (**c**) and hydrogen bond patterns formed by sialic acid in clusters L24 (**b**) and L25 (**d**). In (**a**,**c**) Protein chains are superimposed by the carbohydrate binding site alignment algorithm. b.Top: Superantigen-like protein 11 from *Staphylococcus aureus* in complex with Sialyl Le^x^. Bottom: Putative pertussis-like toxin in complex with α2,6-SiaLacNac from *Salmonella enterica*. The β-sheet region binding sialic acid is composed of a Serine/Threonine interacting via its side chains with the carboxylic acid moiety, followed by two aromatic amino acids, which further stabilize carbohydrate binding. Three other complexes of cluster L24 are provided in Fig. S11. **d**. Top: α2,3-SiaLac binding site on canine adenovirus type 2 fibre head. Bottom: Sialic acid binding site on paired immunoglobulin-like type 2 receptor beta. Interaction pattern involves a glutamine (interacting with the carboxylic acid moiety) followed by an aromatic residue in the first β-sheet and an arginine from the second β-sheet (interacting with the carboxylic acid moiety). The glycerol group interacts with backbone atoms of either a serine or a glutamine. (**e-k**) Similar mannose binding patterns (cluster L1) found in lectins from different kingdoms of life and different folds. L-type legume lectin from *Macropsychanthus grandiflorus* (**e**), archaea lectin (**f**), natterin like lectin from zebrafish (**h**) and jacalin lectin from banana (**i**) are aligned to representative L1 structure of lectin from *Centrolobium tomentosum* (in yellow). Binding site superposition to plant lectin at residue level is shown for archaea and zebrafish lectins in (**g**), with detailed residue annotation provided in (**k**). Key functional residues highlighted in gray according to the family-specific loop nomenclature. Protein functional annotation and alignment scores are provided in (**j**).

The characteristic binding pattern observed in cluster L24 seems specific for bacterial lectins. In comparison, interaction of sialic acid with animal lectins from cluster L25 (Fig. 2c) also appears in the proximity of β-sheets. However, the carbohydrate ring is oriented differently. In addition to the carboxylic acid group, the glycerol group of sialic acid participates in the hydrogen bond formation with the backbone atoms of the β-sheet residues (Fig. 2d).

#### Similar binding patterns in mannose/glucose binding lectins of different architectures and UniLectin classes

While sialic acid binding sites are conditioned by its particular chemical composition, simple carbohydrates such as glucose and mannose can also form characteristic binding sites found in lectins of divergent folds and coming from different organisms. For instance, cluster L1 contains both mannose and glucose binding sites found in lectins from different UniLectin classes found in animals, plants and archaea (Fig. 2e-k, Tab. S5). For the legume (Fig. 2e) and archaea lectins (Fig. 2f) of the same b-prism fold we obtain an almost perfect superposition of the five loops (called A to E, Fig. **2g top**) known to confer carbohydrate specificity^40,41^. Moreover, avery similar binding site is detected in zebra-fish Natterin-like and banana Jacalin-like lectins of different topologies (Fig. 2h,i) formed by conserved GG- and binding loops, and a variable recognition loop granting lectin specificity^42^. Indeed, we can establish a clear correspondence between the Jacalin-like GG loop and legume lectin loop B, between the binding loop and loop D, while the recognition loop partially aligns with loop C (Fig. 2g, bottom).

A closer look at the atomistic details of binding site composition further highlights CBS similarity at the level of the conserved loops (Fig. 2k). Indeed, GG- and B-loop match perfectly (Fig. 2k, in green). We observed a nearly perfect match between aspartic acid and glycine found in binding or D-loops (Fig. 2k, in blue). Furthermore, the conserved GXXXD motif of the Jacalin lectin binding loop (131-135 in 4ZNQ) crucial for primary carbohydrate binding^42^ is partially replicated in L-type lectins (5EYX) with one glycine residue from loop D and an asparagine residue from loop A.

### Filling in missing annotations of carbohydrate-binding proteins

Carbohydrate binding site clustering allowed us to efficiently explore the important number of binding sites with no consistent annotation in the existing databases^13,19,20^. By explicitly comparing unannotated CBS to representative binding site structures, we have detected highly resembling binding sites for 621 binding sites. Among them, 241 CAZy active sites are already attributed a correct CAZy annotation but lack active site annotation of UniProt. At the same time, we detect at least six different proteins bearing CBM motifs, confirmed by literature analysis (Tab. S6) but not referenced in CAZy. Several of these proteins correspond to SusE/SusF, known for their involvement in starch utilization systems in the human gut microbiota^43^. These proteins exhibit a significant level of structural similarity to known CBMs but show a low level of sequence conservation, being one of the fastest-evolving among Polysaccharide Utilization Loci^44^. This makes traditional sequence-based methods, such as BLAST or HMM, insufficient for their identification and highlights the necessity of structure-based approaches to reliably capture these fast-evolving proteins.

Finally, our algorithm detects 20 proteins with unique UniProt IDs and no annotation in UniLectin as bringing lectin binding sites on their surface (Tab. S7). To verify our mapping results, we have compared binding site alignment scores with the scores provided by the reference tool for lectin identification from protein sequence, LectomeXplore^14^. We have compared scores for both the proteins referenced in UniLectin and for unannotated proteins (Fig. 3a,b). Interestingly, the observed correlation is rather low

**Figure 3.**
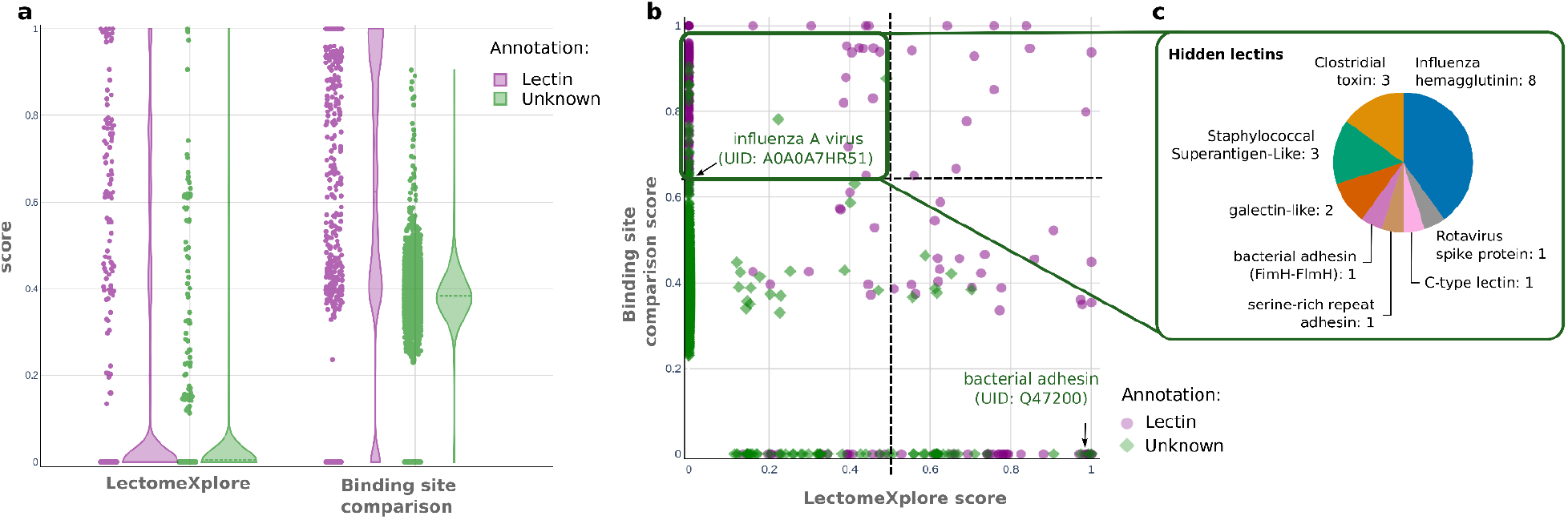
Comparison of sequence-based method for lectin annotation (LectomeXplore) with the results of local binding site alignment. Each point corresponds to a Uniprot ID for which at least one of the following conditions is satisfied: LectomeXplore score is greater or equal to zero; or the highest score of binding site superposition is obtained for a lectin cluster. Violin plots of scores obtained by two methods (**a**) accompanied by a scatter plot (**b**). Violet color stands for known lectins (references in UniLectin3D) and green color marks previously unannotated proteins. In panel (**c**) we provide proportions of the different UniLectin classes predicted by structure superposition score (>0.65) with low LectomeXplore score (<0.5).

(Fig. 3b) and there are very few lectins with high LectomeXplore and structural alignment scores (Fig. 3b, top right). Moreover, for known lectins (Fig. 3a, in purple), LectomeXplore scores remain rather low and only 16% is above the 0.5 threshold. In comparison, the binding site superposition scores do not fall below 0.38 and reach 0.65 and above (our threshold) for 50% of proteins (Fig. 3a). Nevertheless, the sequence-based score appears more precise than our method for several examples (Fig. 3b, bottom right) such as bacterial adhesin binding N-acetyl-glucosamine correctly predicted as lectin by LectomeXplore despite its missing annotation in UniLectin and low binding site superposition score (Fig. 3b, highlighted).

For proteins with LectomeXplore score above zero and high binding site comparison score both methods predict the same UniLectin class except for one example: K4LM89 which we successfully predict as a Caliciviridae (Norovirus/Lagovirus) VP capsid while LectomeXplore predicts as a FMDV receptor complex with a high confidence (0.76) (Tab. S8).

All the 20 “hidden lectins” have LectomeXplore score below 0.5 and it is equal to zero for 18 of them (Fig. 2p, top left; Tab. S7). Nevertheless, according to their functional annotations, most of these proteins can be classified as lectins. Most of them are associated with viral or bacterial infections (Fig. 3c) and high sequence divergence prevents their correct identification by the sequence-based methods. Nevertheless, CBS on the protein surface is often well conserved and detected by our approach, as it is the case for the sialic acid binding site of haemagglutinin H10 from the H10N7 strain of influenza A (Fig. **3c**).

## DISCUSSION

Despite their abundance and importance, the analysis of protein-carbohydrate binding interfaces remains particularly challenging due to significant variability and issues related to their experimental resolution. Indeed, according to our results, more than half of different carbohydrate binding sites resolved in the PDB are not annotated as such due to the particularities of ligand annotations. This under-representation suggests a broader diversity of carbohydrate interactions than was previously acknowledged.

The absolute majority of the currently available annotation tools are based exclusively on protein sequence analysis without considering the crucial role of local amino acid arrangement often responsible for carbohydrate recognition. Our findings based on thorough structural comparison suggest that despite high diversity of carbohydrate binding site structures, some binding patterns can emerge in proteins with no detectable homology relationship due to the local evolutionary convergence. The phenomenon of convergent evolution in recognition patterns of binding sites has been previously documented^45,46^, and underscores the complexity and adaptability of protein functions across diverse biological systems. Therefore, our insights bring particularly important information when studying the specificity of carbohydrate binding for different protein classes, which is necessary for the development of new therapeutic strategies. For example, the striking resemblance between sialic acid binding sites found in different bacteria suggest a potential for the development of a broad-spectrum peptide-inhibitors, which would target adhesion proteins by replacing sialic acid in the β-sheet pattern formation.

In conclusion, multi-scale analysis of the diversity of protein-carbohydrate interfaces highlights the intrinsic complexity of these interactions. While the presented results allow us to complement the available knowledge on several classes of carbohydrate-binding proteins, such as carbohydrate-binding enzymes, lectins and carbohydrate binding modules, the applied protocol can be further adapted to such important categories as sugar transporters or glycosaminoglycan binding proteins. In particular, while the present study is focused on binding site similarities identified for individual carbohydrate moieties, a more relevant biological unit for such classes of interaction as GAG recognition would be an oligosaccharide fragment. However, such methodological modification would require systematic validation of the cumulated alignment scores, which would certainly be possible soon following the growing amount of available experimentally resolved complexes and general information on GAG recognition^47^. Nevertheless, in the current form our algorithm already allows successful identification of similar patterns formed by oligosaccharide ligands of different size and provides an important contribution for functional annotation of carbohydrate binding proteins, complementing the information available from protein sequence analysis.

The reported results open a new perspective to the investigation of specificity and mechanism of action of carbohydrate-binding proteins and have potential to contribute to sugar-related drug development.

## ACKNOWLEDGMENTS

The study was performed in the framework of the SugarPred project funded by the French National Research Agency [grant no. ANR-21-CE45-0019] with the support of the Data Intelligence Institute of Paris (DiiP). A part of calculations was performed using high performance computing resources of TGCC (Très Grand Centre de Calcul) [grant no. A0140714131] funded by GENCI (Grand Equipement National de Calcul Intensif, France) as well as the IFB-core cluster (Institut Français de Bioinformatique, France). The authors also thank the Ministry of Research (France), Université Paris Cité (France), National Institute for Health and Medical Research (INSERM, France) and IdEx [grant no. ANR-18-IDEX-0001].

## AUTHOR CONTRIBUTIONS

Original study design: TG. Data extraction & analysis: AG, TG. Methodological development: AG, FG, TG. Interpretation of the results: AG, SP, TG. Original draft: AG, TG. Manuscript revision and final draft: AG, FG, SP, TG.

## DATA AVAILABILITY

All the data analyzed in the current study are available at https://www.dsimb.inserm.fr/DIONYSUS/

