## Supplementary Data for "Unraveling the diversity of protein-carbohydrate interfaces: insights from a multi-scale study"

**DIONYSUS database content** **2**
Carbohydrate definition according to Protein Data Bank 2
Diversity of carbohydrate-bringing ligands 2
Diversity of carbohydrate binding sites for different proteins 3
**Diversity of protein-carbohydrate interfaces formed by identical proteins** **5**
**Similarity of protein-carbohydrate interfaces among proteins from different classes** **7**
CAZy active sites 7
Carbohydrate binding modules 10
Lectin binding sites 12
**Mapping of unannotated CBS** **13**

### DIONYSUS database content

#### Carbohydrate definition according to Protein Data Bank

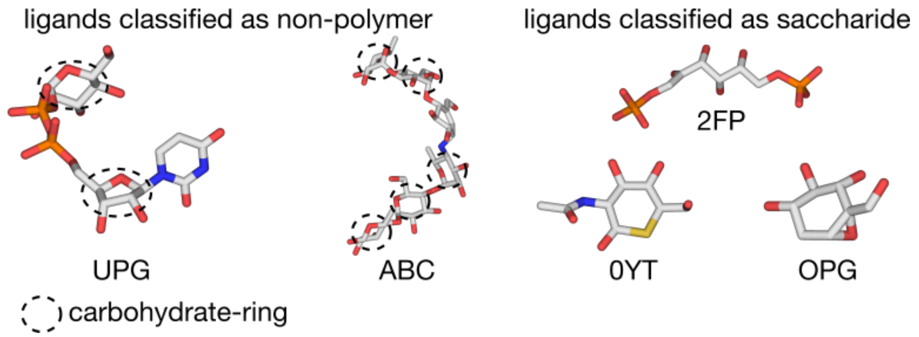


**Figure S1.** (*left*) 3D structures of ligands classified as non-polymer by PDB annotations containing multiple carbohydrate rings (circled in dotted lines). (*right*) 3D structures of ligands classified as saccharide by the PDB or ProcarbDB, but not containing a conventional carbohydrate ring.

#### Diversity of carbohydrate-bringing ligands

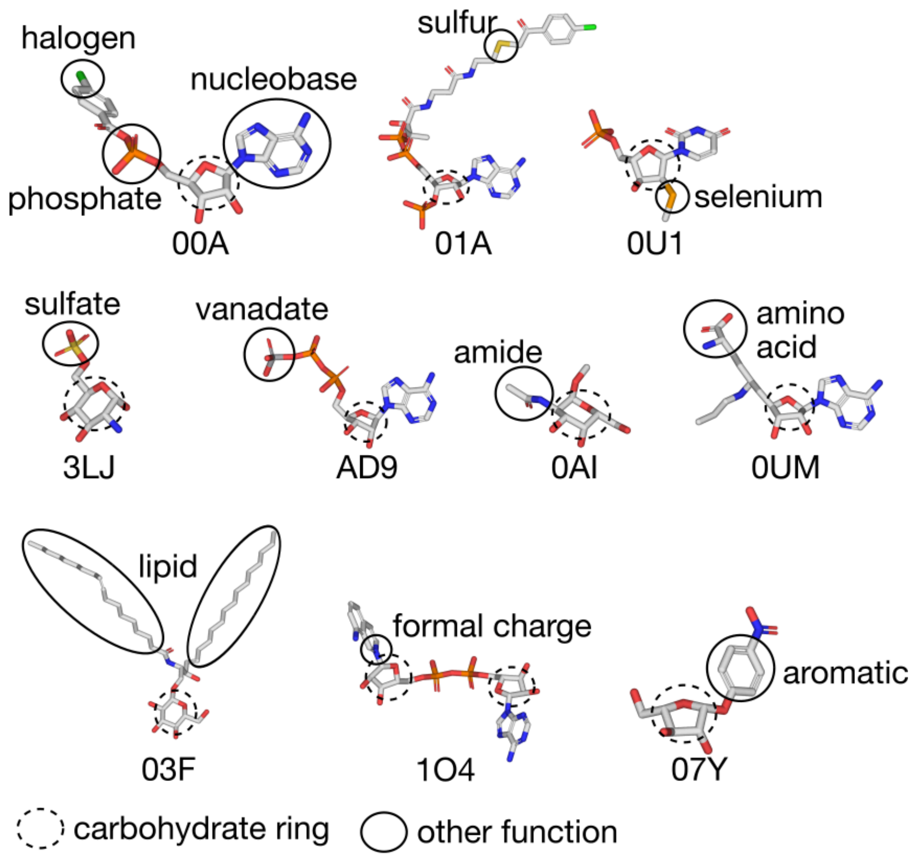


**Figure S2.** 3D structures of carbohydrates with additional chemical functions.

#### Diversity of carbohydrate binding sites for different proteins

*Lectin binding sites.* In accordance with UniLectin, the majority of detected carbohydrate-protein interfaces are crystallized for galectin-like lectins (Fig. S4), followed by numerous mannose-specific lectins: L-type legume lectins, Jacalin-like lectin, Monocot-lectin like and fucose-specific lectins: AAL-like. We also report a variety of lectin binding sites found in viruses (such as influenza hemagglutinin and rotavirus spike protein) and in bacteria (e.g., Cholera toxin like AB5). In terms of the general fold, as expected, most of the extracted structures consist of β-sheets (β-sandwich being the most common, Fig. S5) while lectins of hybrid α/β OB-fold account only for 7% of lectin sites.

*Enzyme binding sites*. The most represented enzyme family responsible for 20% of carbohydrate binding site diversity among enzymes is GH13 (Fig. S4) acting on all α-glucoside linkages. The GH13 family gathers both enzymes with various enzymatic activities (the most common being α-amylase) and proteins with lost glycosidase activity during evolution[1]. The rest of most common families together represent less than 6% of the dataset formed by CBS from GH10 (xylanases), GH6 (cellulases) and GH9 (cellulases).

*CBM binding sites*. CBM48 and CBM20 are responsible for 30% of CBS diversity among carbohydrate binding modules (Fig. S4), followed by CBM13 and CBM35. Both families, CBM48 and CBM20, are considered as starch binding domains[2] and are often found in close interaction with GH13 enzyme domains. The diversity in binding sites formed by CBM48 and CBM20 at the level of one carbohydrate residue comes from their ability to accommodate different arrangements of linear or cyclic glucans with low specificity for their composition. Similarly, both CBM13 and CBM35 families gather modules participating in xylan polysaccharide binding[3]. Interestingly, CBM13 constitutes in fact a limit-case because numerous proteins containing this module are considered lectins, which lead to the creation of the UniLectin family CBM13-Xylanase. For the sake of consistency, we always considered such sites as CBMs.

*Antibody binding sites.* Among antibodies, the small amount of data is hard to interpret because of the small sample size, but it appears that more binding sites are formed with an antibody heavy chain than with an antibody light chain. Furthermore, among heavy chains the most common subclass is IGHV1 and among light chains, IGKV1.

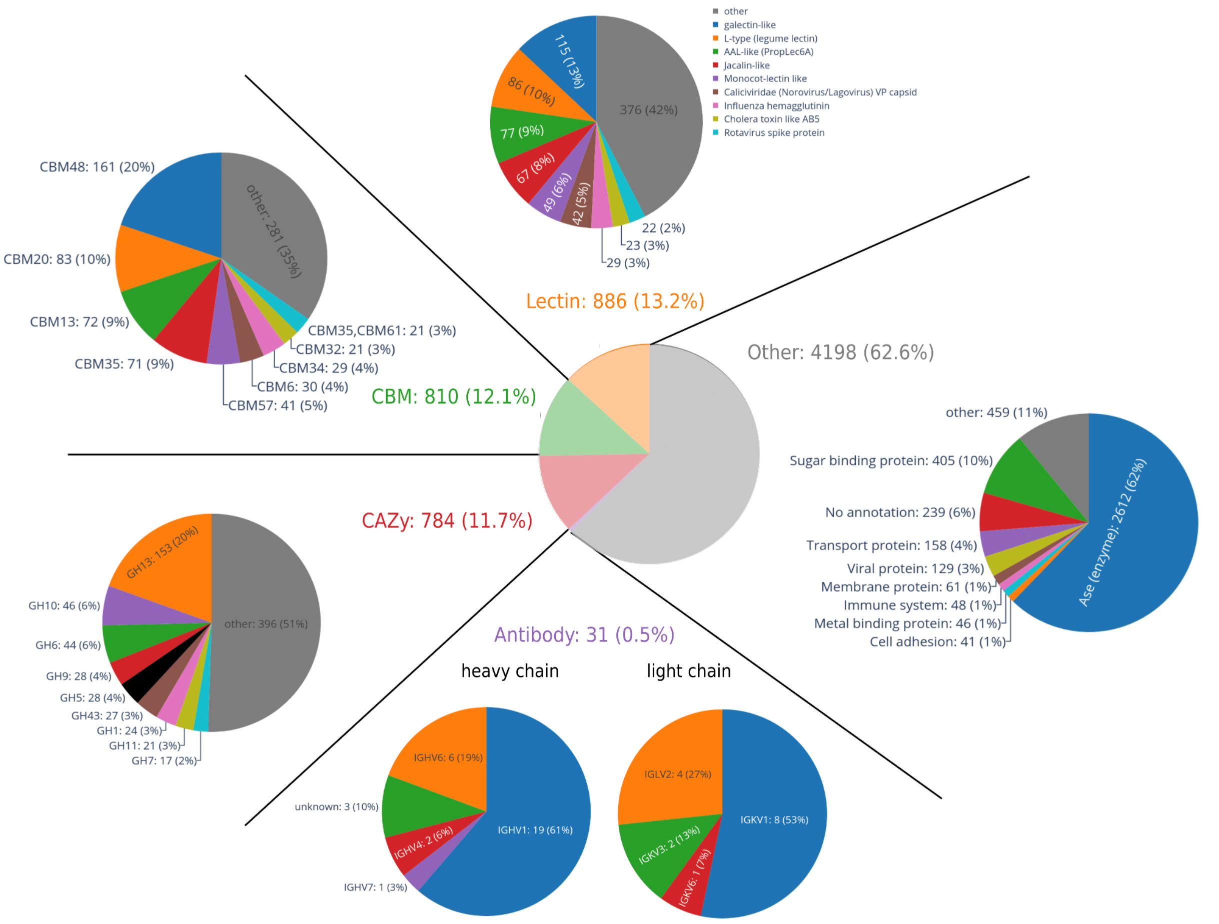


**Figure S3.** Proportion of unique carbohydrate binding sites (see Materials & Methods for the details) from different protein classes according to existing databases.

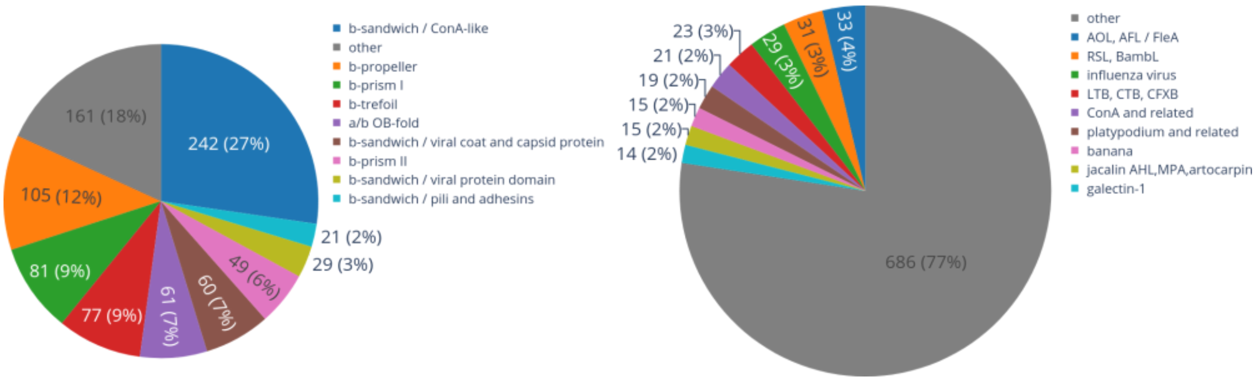


**Figure S4**. Pie chart representing the proportion of the top 10 UniLectin folds (left) and families (right)

#### Diversity of protein-carbohydrate interfaces formed by identical proteins

High sequence identity, even when combined with the same carbohydrate name, does not necessarily confirm that binding sites are identical between two proteins. Conversely, low sequence identity or different ligand names do not prevent the possibility of nearly identical carbohydrate binding sites (CBS) on a protein surface. In Fig. S6 we report several examples of identical proteins forming different interfaces with the same carbohydrate residue and discuss the underlying biological phenomena.

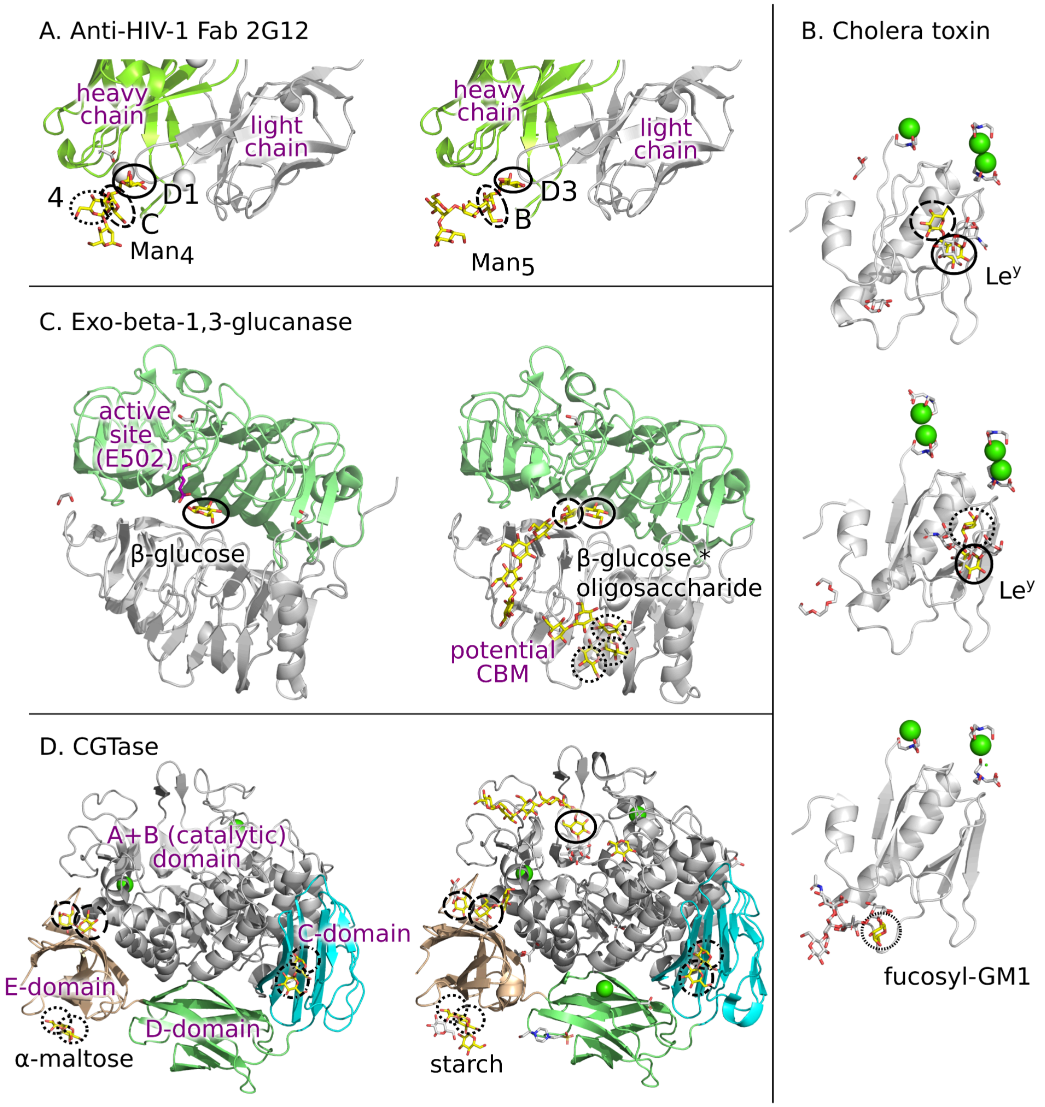


**Figure S5.** Localization and size of redundant groups found for some interesting sites in each category. Carbohydrates of interest are coloured in yellow. **A**. Anti-HIV-1 Fab 2G12 (PDB IDs: 6MSY and 6MUB) bound to mannose. Heavy chain coloured in gray and light chain coloured in green. **B**. Cholera toxin monomer (PDB IDs: 5ELB, 5ELF, 6HMY) bound to fucose-containing $Le^y$ and fucosyl-GM1. **C.** Exo-beta-1,3-glucanase bound to β-glucose oligomer (PDB IDs: 4TYV, 4TZ5) with the active site reverse mutated from alanine to glutamic acid to show the localization of the active site. Sites classified as CAZys are circled with dashed lines, while sites classified as “other” are circled in dotted lines. **D.** CGTase (PDB IDs: 1CDG, 1OT2) bound to α-glucose.

*Mannose binding patterns of $Man_9GlcNAc_2$ on the 2G12 surface.* In Fig. S6A we report two structures of a human monoclonal antibody 2G12, for which we identify three different mannose binding patterns among nine resolved complexes with different α-mannose oligosaccharides. Specificity of Fab 2G12 was in the center of numerous studies due to its ability to act against the human immunodeficiency virus type-1 (HIV-1) through its interaction with a dense cluster of oligomannoside on the surface of the gp120 envelope protein of HIV-1[33,34]. In particular, recent findings show that 2G12 can bind D1 and D3 arms of $Man_9GlcNAc_2$ with broad specificity[35]. In accordance with these studies, our analysis identifies as almost identical D1 and D3 mannose binding sites (Fig. S6A, bold circles), as well as binding sites B and C (Fig. S6A, dashed circles), with most superficial residue 4 forming a different pattern.

*Different fucose binding modes of cholera toxin.* Cholera toxin is known to target both monosialotetrahexosylganglioside (GM1) and fucosylated receptors to enter human cells[36–38] leading to different development of the disease in individuals with different blood-type and secretor status. B subunit of cholera toxin (responsible for fucose binding) was crystallized in 7 structures, and due to its pentameric structure, we identified more than 109 monomer binding sites of α-fucose residue. Among all these CBS, we identify only five different binding patterns (Fig. S6B). Four of them correspond to those reported in the literature using $Le^y$ as a model for fucosylated receptors[39], while the last one corresponds to fucosyl-GM1 binding site. Interestingly, our approach robustly detects fucose high affinity binding sites (Fig. S6B, bold circles). At the same time, the other three binding patterns correspond to different binding modes of the same residue (Fig. S6B, dashed and dotted lines in the top-two panels), which are not distinguished in previous studies[39] despite ligand flexibility.

*Glucose binding by exo-β-1,3-glucanase.* The active site of sacteLam55A from *Streptomyces* sp. SirexAA-E, an exo-β-1,3-glucanase from family GH55, is formed by the catalytic acid E502, which breaks β(1-3) bond in the substrate glucose polymer. Among all the available NMR models for 8 available structures, we detect 235 potential enzymatic binding sites. All of them correspond to one of two binding patterns shown in Fig. S6C, allowing the enzyme to fixate the substrate and break the glycosidic bond between two residues. Interestingly, the same binding site always accommodates the terminal residue of the polysaccharide (in accordance with the "exo" characteristic of the enzyme), and is also more frequently found in different structures due to its ability to bind β-glucose monomers (Fig. S6C, left).

*Potential CBM identified in the structure of exo-β-1,3-glucanase.* According to CAZy annotations, the only carbohydrate binding site of exo-β-1,3-glucanase corresponds to its active site analyzed above. Nevertheless, we detect 550 unannotated CBS, corresponding to 11 different binding patterns, among which three binding patterns are located far from the active site and belong to a different domain (Fig. S6C, gray domain). Interestingly, the domain decomposition of this enzyme is inconsistent in ECOD (e.g., in structure 4TYV chain A is split into two domains and identical chain B is considered a monodomain). Therefore, we used SWORD2[40] optimal partition indicating two domains as coloured in Fig. S6C. The distinct structure of the considered domain and CBS location are in accordance with the previously expressed hypothesis that exo-β-1,3-glucanase secondary binding site acts as a carbohydrate binding module[41] and therefore our analysis provides information complementing annotations available in the existing databases.

*Different carbohydrate binding sites in CBMs.* The CAZy database contains a notable example involving cyclomaltodextrin glucanotransferase (GH13_2, CBM20) bound to α-glucose, as shown in Figure S6D.

This enzyme, appearing in 23 structures, is generally divided into four domains, with domains A and B generally fused in a single catalytic domain, and domains C and E known for starch binding[42,43]. Out of 178 binding sites, 135 are reduced to only six non-redundant sites corresponding to three disaccharide poses (two on the E-domain and one on the C-domain) reported by the authors[43]. The remaining nine non-redundant sites correspond to the portions of starch forming weaker contact with the protein.

The reported examples highlight the importance of CBS comparison using residue-wise binding patterns. Indeed, such an approach allows us to detect similarity between ligands of different shapes (as for D1 and D3 arms of $Man_9GlcNAc_2$ in Fig. S6A) or polymers of different length (as for maltose and starch polymer in Fig. S6D).

### Similarity of protein-carbohydrate interfaces among proteins from different classes

#### CAZy active sites

**Table S1**. Annotation and alignment metrics of clusters containing binding sites with different CAZy family annotations

| | Annotations | | | | Organism | Alignment metrics | | | | Catalytic domain residue id | |
|---|---|---|---|---|---|---|---|---|---|---|---|
| Cluster | PDB Entry | CAZy family | CAZy clan | ECOD id | | RM SD | TM-Score | Sequence Identity | Number of equivalent residues | Start | End |
| D3 | 4D47.G | GH68 | GH-J | 5.1.2.7 | *Erwinia amylovora* | - | - | - | - | 4 | 414 |
| | 3RWK.A | GH32 | GH-J | 5.1.2.5 | *Aspergillus ficuum* | 3.26 | 0.61 | 12% | 253 | 24 | 357 |
| | 1W2T.C | GH33 | GH-J | 5.1.2.3 | *Thermotoga maritima* | 3.16 | 0.59 | 13% | 254 | 1 | 294 |
| D53 | 1L2A.A | GH48 | GH-M | 5.1.2.3 | *Clostridium thermocellum* | - | - | - | - | 32 | 663 |
| | 7F82.B | GH8 | GH-M | 5.1.2.5 | *Escherichia coli* | 3.33 | 0.43 | 12% | 273 | 22 | 359 |
| D62 | 6DDU.A | GH2 | GH-A | 10.1.2.234 | *Mus musculus* | - | - | - | - | 324 | 690 |
| | 1UMZ.A | GH16 | GH-B | 10.1.1.14 | *Populus tremula* | 6.23 | 0.19 | 7% | 58 | 6 | 272 |

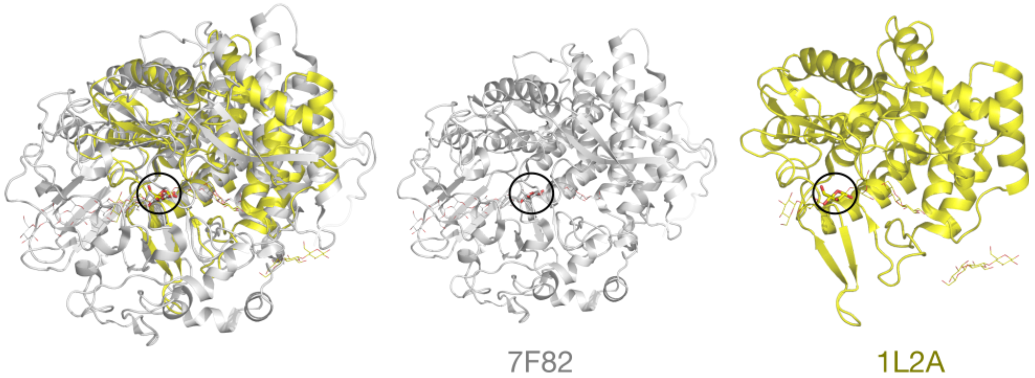


**Figure S6**. Superposition of protein chains forming binding sites from cluster D53. Catalytic domains highlighted in opaque, non-catalytic domains in transparent. Clustered carbohydrates highlighted in sticks, other carbohydrates in lines.

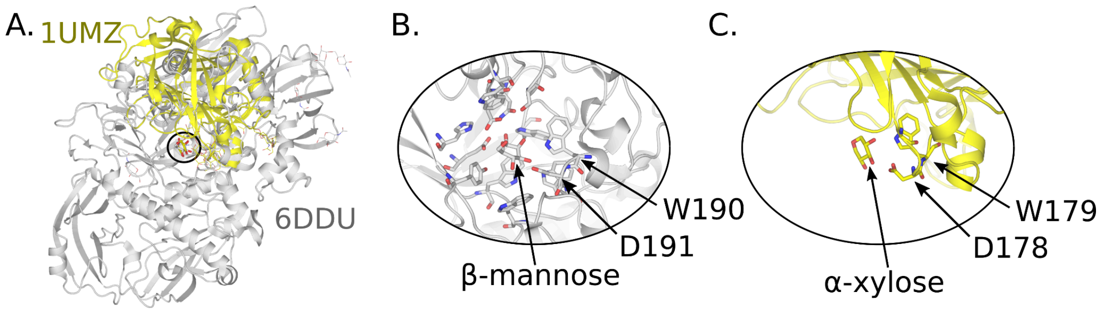


**Figure S7.** Superposition of protein chains establishing binding sites in cluster D62 with (A) and their binding sites (B,C). The correspond to different clans and share no detectable homology at the level of general fold: β-mannosidase from clan GH-A and a Xyloglucan endotransglycosylase from clan GH-B. At the same time, they share the same retaining mechanism and use glutamic acid as catalytic nucleophile and proton donor in the active site.

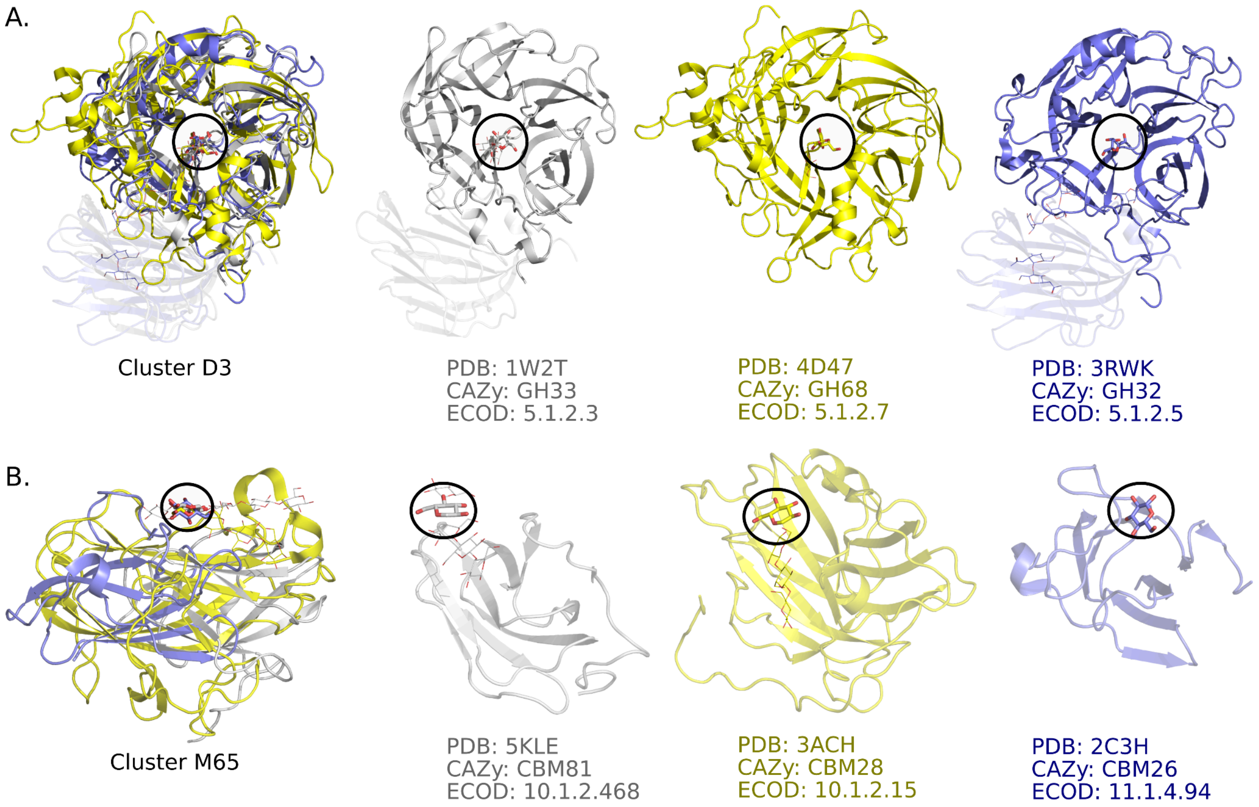


**Figure S8.** Proteins with divergent sequences and folds, sharing similar structure of carbohydrate binding sites. **A**. Similar fructose binding sites found in glycoside hydrolases with low sequence identity (<15%). **B**. Similar glucose binding sites found in CBM from different families and folds.

#### Carbohydrate binding modules

**Table S2 (separate file).** Semi-manual attribution of CBM family to each clustered domain. We used both annotations to identify CBM domains, by excluding those with catalytic annotations. The attributed CBMs are reported in Table S2. The peculiar location of CBM on non-catalytic domains which are defined ambiguously makes their automatic identification using CAZy and ECOD annotation impossible. For instance, we find that some clusters, such as M12, mainly have sites on domains identified as catalytic by ECOD (Alpha-amylase), even though the corresponding sites lack any active site annotation. Other domains such as M35 show shared binding between the catalytic and CBM domains.

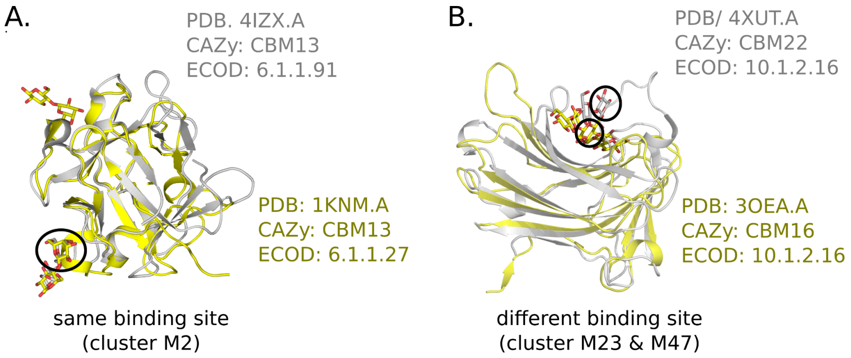


**Figure S9**. ECOD family similarity vs carbohydrate binding site similarity. **A**. Superposition of CBS found in protein chains with different ECOD IDs (with same CAZy annotation) but attributed to the same cluster of binding sites (M2). **B**. Superposition of two CBM coming from the same ECOD family but having different annotation according to CAZy and binding sites belonging to different clusters.

**Table S3**. Characteristics of clusters containing different CBMs.

| Cluster | N° sites | N° proteins | Carbohydrates | CBM classes |
|---|---|---|---|---|
| M11 | 3 | 3 | GLC (100%) | CBM41,CBM58 |
| M13 | 5 | 4 | BMA (60%); BGC (20%); GLC (20%) | CBM27,CBM80 |
| M65 | 3 | 2 | GLC (100%) | CBM21,CBM68 |
| M27 | 2 | 2 | BMA (100%) | CBM27,CBM80 |
| M50 | 3 | 2 | XYP (67%); BGC (33%) | CBM15,CBM81 |
| M65 | 3 | 3 | BGC (67%); GLC (33%) | CBM26,CBM28,CBM81 |

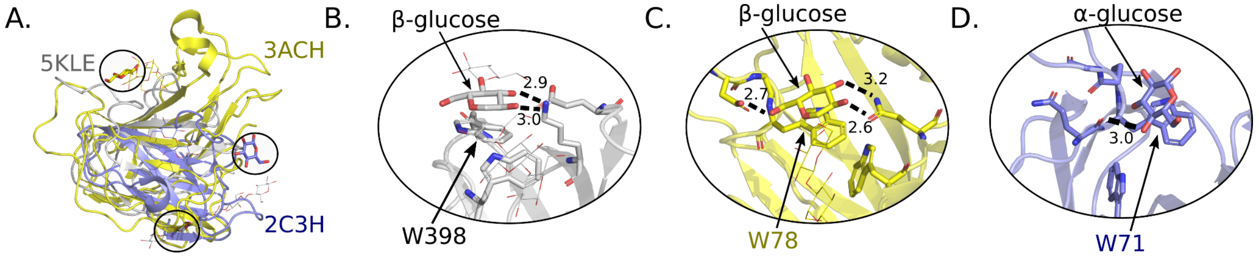


**Figure S10**. Glucose binding sites from cluster M65. **A**. Global alignment using TM-align. **B-D**. Detailed structure of binding sites found in chains 5KLE.A, 3ACH.A and 2C3H.D respectively.

**Table S4.** Annotations and alignment metrics of the CBM mentioned in Figures S10 and S11.

| | Annotations | | | Alignment metrics | | | |
|---|---|---|---|---|---|---|---|
| Cluster | PDB Entry | CAZy family | ECOD id | RMSD | TM-Score | Sequence Identity | Equivalent residues |
| M2 | 4IZX.A | CBM13 | 6.1.1.91 | - | - | - | - |
| | 1KNM.A | CBM13 | 6.1.1.27 | 1.78 | 0.78 | 24% | 89% |
| M23 | 3OEA.A | CBM16 | 10.1.2.16 | - | - | - | - |
| M47 | 4XUT.A | CBM22 | 10.1.2.16 | 2.93 | 0.73 | 18% | 69% |
| M65 | 3ACH.A | CBM28 | 10.1.2.15 | - | - | - | - |
| | 5KLE.A | CBM81 | 10.1.2.468 | 3.49 | 0.34 | 6% | 73% |
| | 2C3H.D | CBM26 | 11.1.4.94 | 4.03 | 0.24 | 5% | 52% |

#### Lectin binding sites

**Table S5**. Characteristics of lectin clusters gathering proteins from different UniLectin classes. The other clusters generally involve two or three sites with specificity for the same monosaccharide such as β-galactose-binding sites (clusters L32, L66, and L6), α-mannose-binding sites (cluster L47) and α-fucose-binding sites (cluster L27), which demonstrate notable similarity between Fungal prism lectins and Norovirus capsids.

| Cluster | N° sites | N° proteins | Carbohydrates | Unilectin classes |
|---|---|---|---|---|
| L1 | 40 | 20 | MAN (38%), MMA (22%), GLC (18%), GYP (10%), BGC (5%), BMA (5%), AMG (2%) | L-type (legume lectin) (65%), Jacalin-like (28%), Natterin-like (5%), ArchLec-like (2%) |
| L2 | 45 | 28 | GAL (80%), SGA (9%), 1GN (4%), GLA (2%), SDY (2%), GLC (2%) | galectin-like (98%), Turkey siadenovirus A (2%) |
| L4 | 40 | 21 | BGC (48%), GLC (38%), GAL (8%), 6S2 (2%), FRU (2%), FUC (2%) | galectin-like (95%), Turkey siadenovirus A (5%) |
| L24 | 9 | 8 | SIA (89%), NGC (11%) | serine-rich repeat adhesin (44%), Staphylococcal Superantigen-Like (22%), Thyphoid toxin like AB5 (11%), Clostridial toxin,Clostridium HA,Ricin-like (11%), MAR Micronemal protein (11%) |
| L25 | 2 | 2 | SIA (100%) | I-type lectin (50%), Fiber knob (50%) |
| L27 | 3 | 3 | FUC (100%) | Fungal prism lectins (67%), Caliciviridae (Norovirus/Lagovirus) VP capsid (33%) |
| L32 | 5 | 5 | GAL (100%) | Staphylococcal Superantigen-Like (40%), Clostridial toxin,Clostridium HA,Ricin-like (20%), serine-rich repeat adhesin (20%), Thyphoid toxin like AB5 (20%) |
| L45 | 2 | 2 | MAN (50%), GLA (50%) | OAA-like (50%), AAL-like (PropLec6A) (50%) |
| L47 | 2 | 2 | MAN (100%) | L-type (legume lectin) (50%), ERGIC-VIP L-type (50%) |
| L66 | 2 | 2 | GAL (100%) | Ricin-like (50%), Sclerotinia trefoil lectin-like (50%) |
| L67 | 2 | 2 | GAL (100%) | Ricin-like (50%), Earthworm lectin-like (50%) |

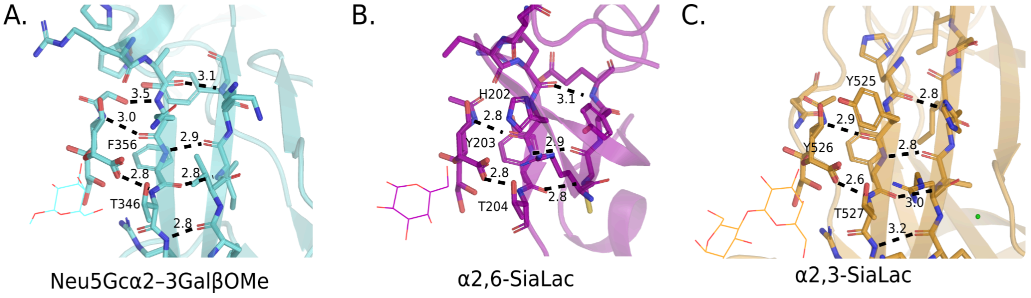


**Figure S11.** Hydrogen bond patterns formed by sialic acid in cluster L24. **A**. SrpA adhesin in complex with Neu5Gc-α2,3-GalβOMe from *S. sanguinis* (PDB ID: 5EQ3, ECOD ID: 11.1.1.2556). **B**. Microneme protein 1 from *T. gondii* in complex with α2,6-SiaLac (PDB ID: 2JH7, ECOD ID: 3083.1.1.1) **C**. HA70 in complex with α2,3-SiaLac from *C. botulinum* (PDB ID: 4LO5, ECOD ID: 10.1.2.63). In the Neu5GC binding site the second aromatic residue is absent, which might be explained by the ability of Neu5Gc to extend the hydrogen bond pattern further with its N-glycolyl moiety.

#### Mapping of unannotated CBS

**Table S6.** Hidden CBM

| Uniprot ID | PDB Id | Carbohydrate | Corresponding CBM | score |
|---|---|---|---|---|
| A0A2N0URA4 | 7RFT | glucose | CBM41 | 0.65 |
| A0A412DXQ2 | 7RPY | glucose | CBM20 | 0.73 |
| G8JZS6 | 4FE9 | glucose | CBM20 | 0.68 |
| G8JZT0 | 4FCH | glucose | CBM20 | 0.66 |
| Q9FEB5 | 4PYH | glucose | CBM20 | 0.79 |
| Q8LK69 | 7VWB | galactose | CBM80 | 0.69 |

**Table S7.** Hidden lectins.

| Uniprot ID | Predicted fold | Predicted class | Predicted family | Score | Score LectomeXplore |
|---|---|---|---|---|---|
| A0A0A7HR51 | b-sandwich / viral protein domain | Influenza hemagglutinin | influenza virus | 0.65 | 0 |
| A0A0R4I961 | b-sandwich / pili and adhesins | bacterial adhesin (FimH-FlmH) | FimH | 0.84 | 0 |
| A0A348FV55 | b-sandwich / viral protein domain | Influenza hemagglutinin | influenza virus | 0.68 | 0 |
| A5H0J8 | b-trefoil | Clostridial toxin | TeNT | 0.68 | 0 |
| C4P282 | b-sandwich / viral protein domain | Influenza hemagglutinin | influenza virus | 0.73 | 0 |
| C9WWY7 | b-trefoil | Clostridial toxin | BoNT/A | 0.86 | 0 |
| G0Z026 | a/b OB-fold | Staphylococcal Superantigen-Like | SSL5 | 0.78 | 0.22 |
| G8IPF0 | b-sandwich / viral protein domain | Influenza hemagglutinin | influenza virus | 0.77 | 0 |
| P08699 | b-sandwich / ConA-like | galectin-like | galectin-3 | 0.86 | 0 |
| P16110 | b-sandwich / ConA-like | galectin-like | galectin-9,galectin-5 | 0.88 | 0.49 |
| P46085 | b-sandwich / Ig-like | serine-rich repeat adhesin | GspB | 0.74 | 0 |
| Q2G1S5 | a/b OB-fold | Staphylococcal Superantigen-Like | SSL5 | 0.82 | 0 |
| Q2G1S8 | a/b OB-fold | Staphylococcal Superantigen-Like | SSL5 | 0.84 | 0 |
| Q3LRX8 | b-trefoil | Clostridial toxin | BoNT/A | 0.9 | 0 |
| Q6DQ34 | b-sandwich / viral protein domain | Influenza hemagglutinin | influenza virus | 0.7 | 0 |
| Q7M462 | a/b mixed / C-type lectin-like | C-type lectin | CD23 | 0.67 | 0 |
| Q91E88 | b-sandwich / ConA-like | Rotavirus spike protein | P[19],Human rotavirus P[6] | 0.89 | 0 |
| R4NN21 | b-sandwich / viral protein domain | Influenza hemagglutinin | influenza virus | 0.71 | 0 |
| V5IRU4 | b-sandwich / viral protein domain | Influenza hemagglutinin | influenza virus | 0.75 | 0 |
| V5IRV0 | b-sandwich / viral protein domain | Influenza hemagglutinin | influenza virus | 0.66 | 0 |

**Table S8**. Comparison between LectomeXplore and our predicted classes on consensus examples

| Uniprot ID | Predicted class | Score | Predicted class by LectomeXplore | Score LectomeXplore |
|---|---|---|---|---|
| A0A090BWT0 | L-rhamnose binding lectin | 0.78 | L-rhamnose binding lectin | 0.69 |
| A0A2K6TQH4 | galectin-like | 0.63 | galectin-like | 0.412 |
| A8MUM7 | galectin-like | 0.61 | galectin-like | 0.4 |
| B0D650 | Laetiporus trefoil lectin-like | 1 | Boletus and Laetiporus b-trefoil lectin | 0.304 |
| C0HK27 | L-type (legume lectin) | 0.82 | L-type legume lectin | 0.385 |
| C1IPK2 | Clostridial toxin | 1 | Clostridial toxin | 0.642 |
| G0Z026 | Staphylococcal Superantigen-Like | 0.78 | Staphylococcal Superantigen-Like | 0.222 |
| K4LM89 | Caliciviridae (Norovirus/Lagovirus) VP capsid | 0.85 | FMDV receptor complex | 0.758 |
| P02870 | L-type (legume lectin) | 1 | L-type legume lectin | 0.159 |
| P04958 | Clostridial toxin | 0.67 | Clostridial toxin | 0.664 |
| P05046 | L-type (legume lectin) | 1 | L-type legume lectin | 0.758 |
| P06750 | Ricin-like | 0.65 | Ricin-like | 0.559 |
| P09382 | galectin-like | 1 | galectin-like | 0.447 |
| P12528 | Salmonella bacteriophage | 0.94 | Salmonella bacteriophage | 1 |
| P16110 | galectin-like | 0.88 | galectin-like | 0.491 |
| P18891 | AAL-like (PropLec6A) | 0.94 | AAL-like PropLec6A | 0.554 |
| P30617 | Monocot-lectin like | 1 | Monocot-lectin like | 0.439 |
| P56217 | galectin-like | 0.94 | galectin-like | 0.405 |
| P58906 | L-type (legume lectin) | 0.95 | L-type legume lectin | 0.392 |
| P81180 | cyanovirin-like | 0.93 | cyanovirin-like | 0.709 |
| P81637 | L-type (legume lectin) | 0.88 | L-type legume lectin | 0.39 |
| Q1W2P6 | galectin-like | 0.95 | galectin-like | 0.458 |
| Q38784 | Monocot-lectin like | 0.95 | Monocot-lectin like | 0.433 |
| Q38789 | Monocot-lectin like | 0.95 | Monocot-lectin like | 0.423 |
| Q8K419 | galectin-like | 0.94 | galectin-like | 0.474 |
| Q8K4Q8 | C-type lectin | 0.72 | C-type lectin | 0.396 |
| Q9H2X3 | C-type lectin | 0.83 | C-type lectin | 0.457 |
| Q9S497 | bacterial adhesin (FimH-FlmH) | 0.8 | bacterial adhesin FimH-FlmH | 0.986 |
| Q9ZFS6 | Staphylococcal Superantigen-Like | 1 | Staphylococcal Superantigen-Like | 0.838 |
| Q9ZKV2 | HOP-OMP adhesins | 0.95 | HOP-OMP adhesins | 0.847 |
